## Supplementary Information (Figures and Tables) for "Quantitative proteome of bacterial periplasmic predation reveals a prey damaging protease"

Lai et al.,

| | |
|---|---|
| Supplementary Figures 1- 4 | page 2 - 5 |
| Supplementary Table 1-3 legends | page 6 |
| Supplementary Tables 1-3 | available online in Supplementary materials |
| Supplementary Tables 4-6 | page 6-9 |
| Supplementary References | page 10 |

### Supplementary Figures

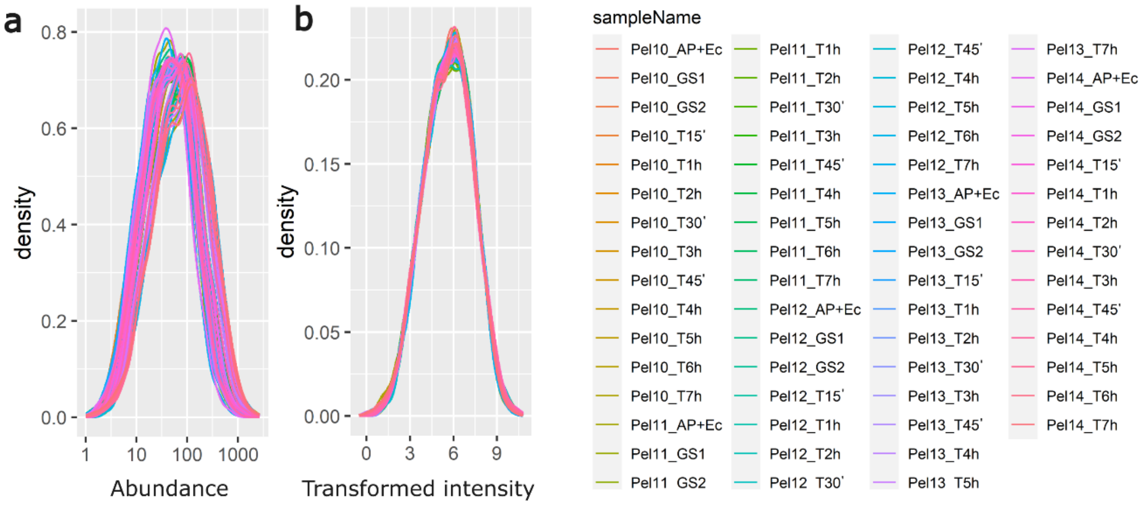


**Supplementary Fig. 1: Density plots showing the effect of sample normalization. a,** Before and **b,** after scaling of protein abundance by internal reference scaling with robust scaling normalizations of all samples of the five biological repeats (Pel10-Pel14). This normalization method takes geometric mean for each protein, calculates scaling factors for each protein to adjust the protein abundance to the geometric mean. The ‘AP+Ec’ condition combines the protein abundances from the ‘attack phase (AP)’ containing *Bdellovibrio* only’ and ‘*E. coli* only (Ec)’, for better comparison with other timepoints. Golden standard samples (GS1, GS2) comprise an equal mixture from each condition and TMT set.

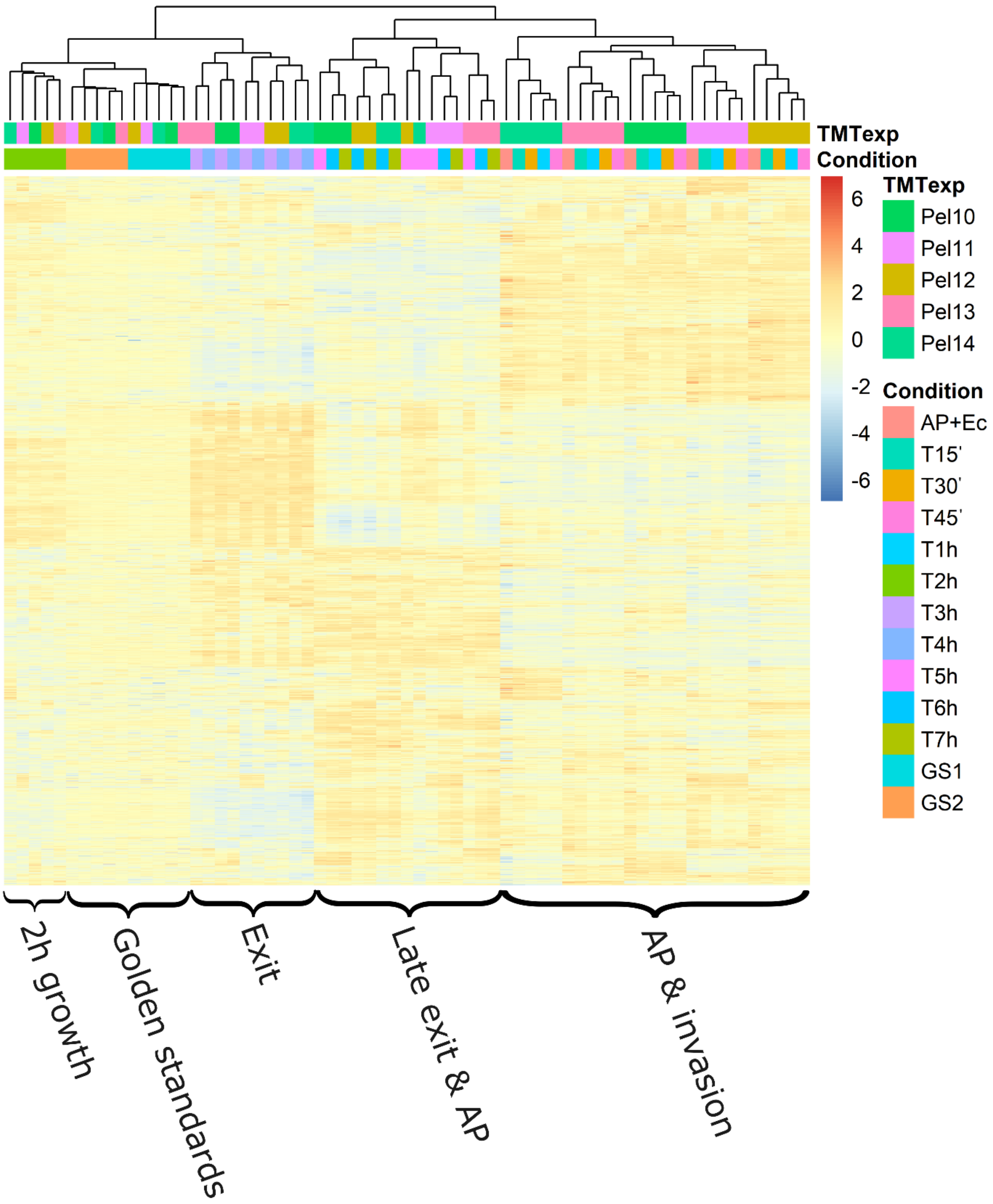


**Supplementary Fig. 2: Hierarchically clustered heatmap showing the relationship between samples with regards to the five biological repeats and the condition.** Proteins were grouped based on the five biological repeats processed in separate TMT sets (TMTexp) and the sampling timepoints of the predatory life cycle (Condition). The 'Condition' timepoints are further grouped categorized by their corresponding life cycle phases, as indicated at the bottom of the heatmap. The units are of relative internal reference scaled intensity of protein abundance from the normalized data. For abbreviations, please refer to Supplementary Figure 1. The golden standard samples (GS1, GS2) cluster together, indicating effective normalization.

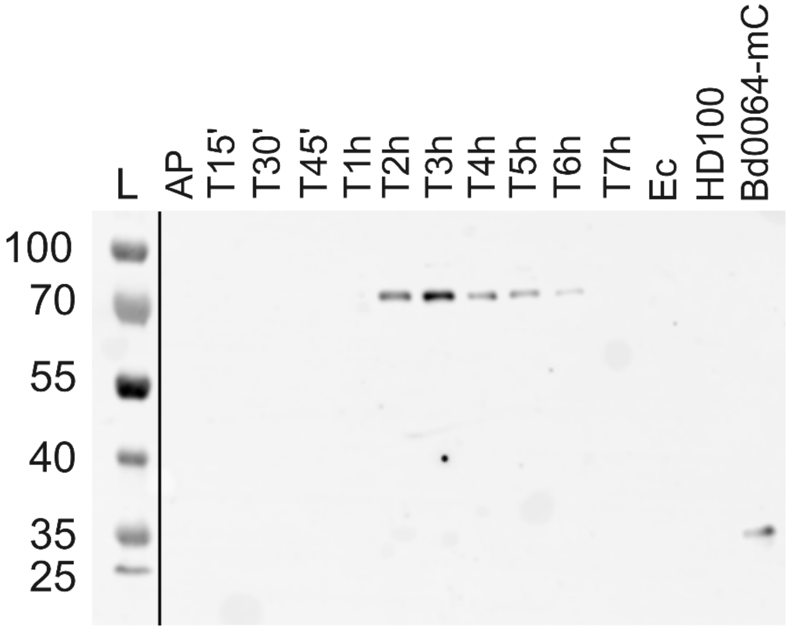


**Supplementary Fig. 3: Western blot of Bd2269-mCherry expression over the whole predatory life cycle.** Semi-quantitative analysis reveals Bd2269 to be most expressed at 3 hours corroborating our quantitative proteome data. The expected size of Bd2269-mCherry is ~83 kDa, while positive control Bd0064-mCherry is ~40.5 kDa. We speculate that the slightly lighter weight band of Bd0064mCherry could be due to partial degradation. AP = 'attack phase' *B. bacteriovorus* HD100 cells; T15'-7h = timepoints in minutes then hours after invasion; Ec = *E. coli* K-12 pZMR100 prey; HD100 = *B. bacteriovorus* HD100 wild-type control; Bd0064-mC = Bd0064-mCherry positive control[1]; L = Protein ladder (kDa) is separated from the Western blot signals by a line to show that it is not chemiluminescent. Two independent biological repeats were performed with the most representative result shown here.

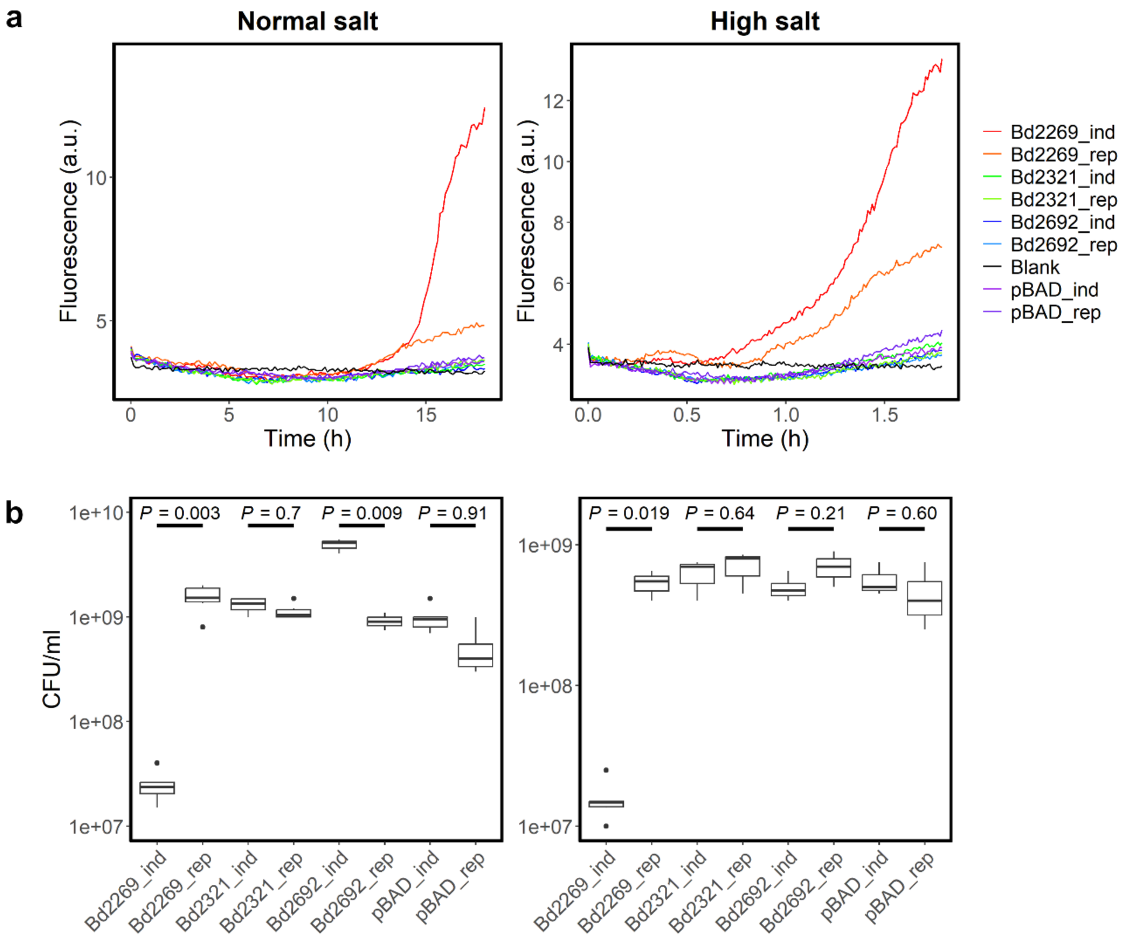


**Supplementary Fig. 4. Second repeat of heterologous expression of protease Bd2269 in *E. coli* S17-1. a,** *E. coli* S17-1 damage from proteases Bd2269, Bd2321 and Bd2692 was tested by β-galactosidase activity assay measuring the fluorescence of chlorophenol red under normal (0.171 M NaCl) and high (0.4 M NaCl) salt conditions over 18 hours. Bd2269, Bd2321 and Bd2692 were heterologously expressed in *E. coli* S17-1 from a pBAD18-based plasmid under inducing ('_ind') or repressing ('_rep') conditions. 'pBAD' stands for the empty vector. *E. coli* S17-1 overexpressing Bd2269 (Bd2269_ind) caused internal damage and β-galactosidase release over time. **b,** Boxplots showing *E. coli* cell viability 18 hours after the start of protease induction ('_ind') or repression ('_rep') in different protease expression strains (Bd2269, Bd2321, Bd2692, pBAD [empty vector control]). Two-tailed Welch two-sample t-tests compare the mean values between the induced and repressed conditions for each protease (*p*-values indicated in graph). Three technical repeats were evaluated across a dilution range of 4. Black dots separate from the box plots represent outliers.

### Supplementary Tables legends:

**(Supplementary Tables 1-3 are available online in Supplementary material as .xls files)**

**Supplementary Table 1: Normalized protein abundances of each protein over a 7-hour period.** Protein abundances are normalized by internal reference scaling with TMM and robust scaling (see methods). Displayed are protein accessions, names, NCBI descriptions, detailed EGGnog descriptions[2], with e-values and the number of unique peptides mapped to each protein.

**Supplementary Table 2: Proteins grouped into the nine clusters of distinct abundance patterns throughout the predatory life cycle.** Each cluster is named as per Fig. 2b. The table includes manually combined cluster numbers and lists the protein accessions for each cluster.

**Supplementary Table 3: A table containing all the $\log_2$ fold change values of *B. bacteriovorus* HD100 proteins between each condition and attack phase.** Displayed are protein accessions, names, NCBI descriptions, detailed EGGnog descriptions[2], with e-values, contrast comparing the different condition to attack phase, $\log_2$ (fold change (FC)), false discovery rate (FDR), the number of unique peptides mapped to each protein, significance ($p < 0.05$) and an indication of whether the protein was experimentally determined or imputed. “-“ = No annotation; #N/A = No e-value. The different contrasts are separated into their respective tabs. The FC values here are used to plot the volcano plots in Fig. 3.

**Supplementary Table 4: Primers used in this study.**

| Primer name | Sequence (5' to 3') | Description |
|---|---|---|
| 2269_EcoRI_F | agcttggaattcatgaaattcaacgtgtttgc | Construction of pBAD18-Bd2269. Addition of EcoRI and SalI restriction sites. |
| 2269_SalI_R | tacgttgtcgacttacttcgcgtcgatcgcag | |
| 2692_EcoRI_F | agcttggaattcatgaaacgtgcattactagt | Construction of pBAD18-Bd2692. Addition of EcoRI and SalI restriction sites. |
| 2692_SalI_R | tacgttgtcgacctactgacagaatggtaggt | |
| 2321_GIB_F | ccatacccgtttttttgggctagcgatgaatcatcttgtgaagggc | Construction of pBAD18-Bd2321 plasmid. Addition of overhangs for Gibson assembly. |
| 2321_GIB_R | acagccaagcttgcatgcctgcaggctatttagccgtataaactagagc | |
| pBAD_F | cacggcagaaaagtccacat | Sequencing of pBAD18-protease constructs. |
| pBAD_R | ctctcatccgccaaaacagc | |
| bd2269KO_up_F | attcacgatacctttcattacaattccccccatatccatgg | Construction of pK18-Δ*bd2269*. Amplification of 1-kb upstream region. |
| bd2269KO_up_R | ggaaacagctatgaccatgattacgtgcagcggctgcgtgagc | |
| bd2269KO_down_F | cgttgtaaaacgacggccagtgccatcttgatcggagcaatcag | |

| | | |
|---|---|---|
| bd2269KO_down_R | aattgtaatgaaaggtatcgtgaatgcggaag | Construction of pK18-Δ*bd2269*. Amplification of 1-kb downstream region. |
| bd2269KO_seq_F | cagcatcccgaccgagataa | Sequencing of pK18-Δ*bd2269*. |
| bd2269KO_seq_R | ggtcattcccgcttacaacg | |
| Bd2269Native-F | cctgcaggtcgactgactgattgacccgctctggtcctaaaatcgaac | Construction pCAT-*bd2269* for complementation. Amplification of *bd2269* and 200bp upstream promoter region. Addition of overhangs for Gibson assembly with pCAT backbone of pFL021. |
| Bd2269Native-R | cgtcgtccttgtagtctcctgctcccttcgcgtcgatcgcagcttg | |
| PCATBackbone-F | taattgactgaagtccactggc | Amplification of pCAT plasmid backbone of pFL021. |
| PCATBackbone-R | cctgcaggcatgcaagcttg | |
| PCAT-F | tgccacctgacgtctaagaa | Sequencing of pCAT-*bd2269* |
| PCAT-R | tggcttaactatgcggcatc | |
| PCAT-M | ggtcttatgcgtctaag | |
| 2269_3p_1kb_F | cgttgtaaaacgacggccagtgccattcactggtgttgctcctaaag | Construction of pK18-Bd2269:mCherry. Amplification of Bd2269 1kb region from 3' end with overlap regions for Gibson assembly. |
| 2269_3p_1kb_R | cttgctcaccatcttcgcgtcgatcgcagc | |
| mCherry_2269_F | gatcgacgcgaagatggtgagcaagggcgag | Construction of pK18-Bd2269:mCherry. Amplification of mCherry with overlap regions for Gibson assembly. |
| mCherry_pK18_R | ggaaacagctatgaccatgattacgctacttgtacagctcgtccatg | |

**Supplementary Table 5: Plasmids used in this study.** Amp$^{R}$ = ampicillin resistance, Kan$^{R}$ = kanamycin resistance, MCS = multiple cloning site.

| Plasmid | Description | Source |
|---|---|---|
| pBAD18 | Arabinose-inducible plasmid (*araBAD* promoter), Kan$^{R}$, used to express *B. bacteriovorus* proteases in *E. coli*. | Gift from Leo Eberl. Guzman *et al.*, 1995[3]. |
| pBAD18-*bd2269* | *bd2269* cloned into MCS of pBAD18 for arabinose induction. | This study |
| pBAD18-*bd2321* | *bd2321* cloned into pBAD18 by Gibson assembly for arabinose induction. | This study |
| pBAD18-*bd2692* | *bd2692* cloned into MCS of pBAD18 for arabinose induction | This study |
| pK18*mobsacB* | Suicide vector (Kan$^{R}$, *lacZα*, *sacB*) used for crossovers into the *B. bacteriovorus* genome. | Gift from Prof. R. E. Sockett. Schäfer *et al.*, 1994[4]. |

| | | |
|---|---|---|
| pK18-Δ*bd2269* | 1-kb upstream and downstream regions of *bd2269* in pK18mobsacB to make bd2269 marker-less gene deletion. | This study |
| pCAT.000 | Self replication level T acceptor vector with *lacZ*. Modified BioBrick vector, from RSF1010: ori, RepB and RepC; $Kan^R$, $Amp^R$, | From Addgene (plasmid #119559). Vasudevan *et al.*, 2019[5]. |
| pFL015 (pCAT-based) | pCAT.000-PmerRNA-RBS-mCherry. Amplification of the pCAT backbone for expression within *B. bacteriovorus*. (Plasmid also contained promotor of merRNA[6], an optimized RBS for B. *bacteriovorus*[7], and an mCherry sequence not directly used in the project. This plasmid was used as template for backbone of pFL021. | Generated by and gift from Florian Lindner. |
| pFL021 (pCAT-based) | pCAT.000-$P_{merRNA}$-RBS-VirF-FLAG. This plasmid is based on pFL015 and was used as template for backbone amplification to include the sequence of a C-terminal FLAG-tag. | Generated by and gift from Florian Lindner. |
| pCAT-*bd2269* | pCAT plasmid containing 196-pb of upstream region of *bd2269* (including natural promoter region), followed by *bd2269* with an additional sequence for a C-terminal FLAG tag (amplified from pFL021). This plasmid was used for complementation of *Δbd2269* to measure exit speed (Fig. 4a). | This study |
| pK18-*bd2269*:*mCherry* | pK18*mobsacB* containing sequence for Bd2269 fused with mCherry at the C-terminus (single crossover) for homologous recombination in *B. bacteriovorus*. Used for Western blot confirmation (Supplementary Fig 4). | This study |

**Supplementary Table 6: Strains used in this study.** $Kan^R$ = kanamycin resistance.

| Strain | Description | Source |
|---|---|---|
| *E. coli* NEB5α | *E. coli* cloning strain (*fhuA2Δ(argF-lacZ)U169 phoA glnV44 Φ80Δ(lacZ)M15 gyrA96 recA1 relA1 endA1 thi-1 hsdR17)* | New England Biolabs (C2987)[8]. |
| *E. coli* S17-1 | *E. coli* strain (*thi, pro, hsdR-, hsdM+, recA*; integrated plasmid RP4- Tc::Mu-Kn::tn) | Gift from Prof. R. E. Sockett. Hanahan, 1983[9]. |
| *E. coli* S17-1 pZMR100 | *E. coli* S17-1 strain containing the plasmid pZMR100 ($Kan^R$), used as prey for kanamycin-resistant *B. bacteriovorus.* | Gift from Prof. R. E. Sockett. Rogers *et al.*, 1986[10]. |
| *E. coli* S17-1 pBAD18 | *E. coli* S17-1 strain used as a negative control for arabinose-protease induction assay containing no protease gene after araBAD promoter | This study |
| *E. coli* S17-1 pBAD18-*bd2269* | *E. coli* S17-1 strain expressing Bd2269 when induced with arabinose. | This study |
| *E. coli* S17-1 pBAD18-*bd2321* | *E. coli* S17-1 strain expressing Bd2321 when induced with arabinose. | This study |

| | | |
|---|---|---|
| *E. coli* S17-1 pBAD18::*bd2692* | *E. coli* S17-1 strain expressing Bd2692 when induced with arabinose. | This study |
| *E. coli* S17-1 pMAL-p2_mCherry | *E. coli* S17-1 strain expressing a maltose binding protein-mCherry fusion with a *malE* signal sequence, directing it to the periplasm. This provides an all fluorescent background to detect division of non-fluorescent *B. bacteriovorus* by fluorescence microscopy. | Gift from Prof. R. E. Sockett. Fenton *et al.*, 2010[11]. |
| *E. coli* K-12 MG1655 | Model *E. coli* strain used as prey for *B. bacteriovorus* predation assays. | Blattner *et al.*, 1997[12]. |
| *E. coli* K-12 MG1655 pZMR100 | *E. coli* K-12 MG1655 strain containing pZMR100 plasmid which confers $Kan^R$. Used as prey for $Kan^R$ *B. bacteriovorus* strains. | This study |
| *B. bacteriovorus* HD100 | *B. bacteriovorus* Type strain, wild-type | Gift from Prof. R. E. Sockett. Rendulic *et al.*, 2004[13]. |
| *B. bacteriovorus* HD100 *bd2269*::*mCherry* | *B. bacteriovorus* HD100 with *bd2269*::*mCherry* integrated into genome by single crossover recombination. Used for Western blots confirmation of Bd2269 protein level. | This study |
| *B. bacteriovorus* HD100 *bd0064:mCherry* | *B. bacteriovorus* HD100 with *bd0064*:*mCherry* integrated into genome by single crossover recombination. Used as a positive control for Western blots confirmation. | Gift from Prof. R. E. Sockett[1]. |
| *B. bacteriovorus* HD100 Δ*bd2269* | *B. bacteriovorus* HD100 with a clean deletion of *bd2269,* used for microscopy exit time analysis. | This study |
| *B. bacteriovorus* HD100 Δ*bd2269* pCAT-*bd2269* | *B. bacteriovorus HD100 Δbd2269* with pCAT-*bd2269* for complementation of *Δbd2269* | This study |
| *B. bacteriovorus* HD100 Δ*bd0314* | *B. bacteriovorus* HD100 with a clean deletion of *bd0314* for microscopy exit time analysis | Gift from Prof. R. E. Sockett. Harding *et al.*, 2020[14]. |
| *B. bacteriovorus* HD100 Δ*bd2269* Δ*bd0314* | *B. bacteriovorus* HD100 with a clean deletion of *bd2269* and *bd0314* for microscopy exit time analysis. | This study |
